# Astrocytic ACSBG1 depletion improves lipid-cytokine signaling and attenuates α-Synuclein pathology in a Parkinson’s disease mouse model

**DOI:** 10.64898/2026.05.20.726454

**Authors:** YoungDoo Kim, Bhupesh Vaidya, Jiyoen Kim, Sara Bitar, Femil Joseph Shajan, Ajay Kumar Verma, Hari Krishna Yalamanchili, Shubham Singh, Huda Y Zoghbi

**Author notes:** These authors contributed equally to this work (YoungDoo Kim, Bhupesh Vaidya).

## Abstract

Astrocytes are key regulators of lipid metabolism, and dysregulated astrocytic lipid processing is implicated in Parkinson’s disease (PD) pathogenesis. Our prior genome-wide screens identified ACSBG1, an astrocyte-enriched acyl-CoA synthetase, as a candidate regulator of α-synuclein (α-Syn) levels. However, how ACSBG1 links lipid reprogramming to inflammatory astrocyte activation and α-Syn pathology remains unknown. We compared the transcriptomic, cytokine, and lipid secretomes of TNF-α and IL-1α stimulated primary astrocytes from wild-type (WT) and *Acsbg1* knockout (KO) mice. In vivo, we crossed Acsbg1 KO mice with a Thy1-α-Syn PD model to assess behavior, neuroinflammation, synaptic integrity, and α-Syn levels. Following cytokine exposure, *Acsbg1* KO astrocytes mounted an attenuated inflammatory transcriptional response, secreting significantly fewer inflammatory mediators (e.g., IL-6, RANTES, MIP-3α) and less long-chain Sphingosine 20:1 than WT astrocytes. Importantly, exogenous Sphingosine 20:1 or cytokines from WT reactive astrocytes induced neuronal α-Syn phosphorylation (pS129). *In vivo, Acsbg1* deletion in Thy1-α-Syn mice reduced astrogliosis, rescued synaptic and behavioral deficits, and decreased total and pS129-α-Syn. These findings establish ACSBG1 as a key regulator of inflammatory astrocyte signaling that contributes to α-Syn phosphorylation via specific cytokine and lipid mediators, identifying ACSBG1 as a novel therapeutic target for modulating astrocyte-neuron communication in PD.

**Graphical abstract:** 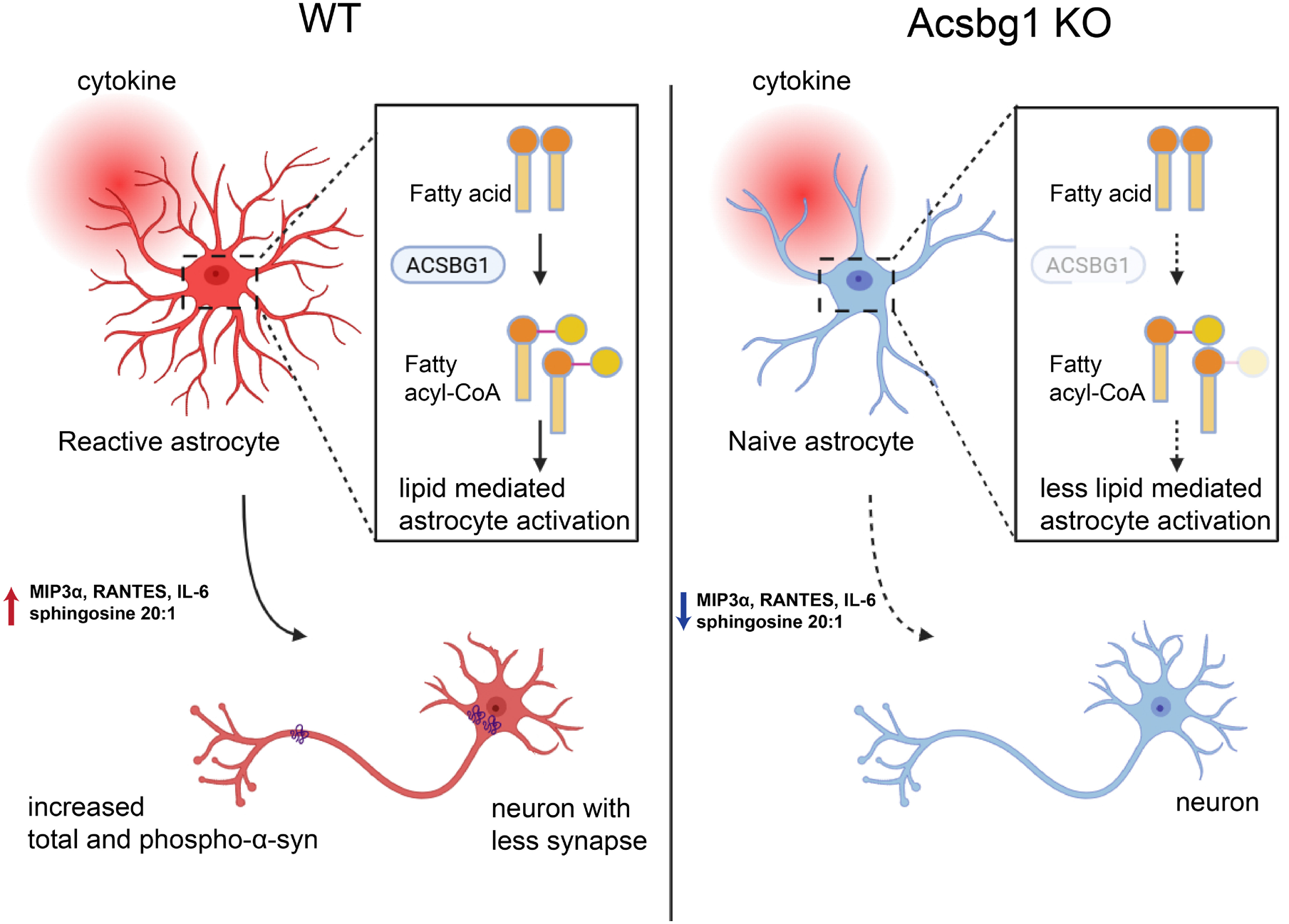

## Introduction

Parkinson’s Disease (PD) affects approximately 10 million individuals worldwide (1, 2), and is primarily identified by motor function deficits such as tremors, rigidity, and bradykinesia. These clinical features result from the progressive degeneration of dopaminergic neurons in the substantia nigra pars compacta (SNpc), and the formation of protein aggregates called Lewy bodies, a hallmark of PD that contain phosphorylated alpha-synuclein (α-Syn) (3).

The *SNCA* gene, which encodes α-Syn protein, was one of the first seven genes identified in familial PD patients (4, 5), and genome-wide association studies have consistently identified *SNCA* as a major risk factor gene for PD (6-8). Total and phosphorylated α-Syn levels are key determinants of PD patients’ symptom severity and disease progression (9).

Consequently, therapeutic strategies aimed at modulating α-Syn levels hold significant promise. Indeed, recent work demonstrated that reducing α-Syn levels by regulating its modulator proteins effectively improves disease phenotypes (10-12). We performed genetic screens to identify genes and pathways that regulate α-Syn levels and identified Acyl-CoA Synthetase Bubblegum Family Member 1 (*ACSBG1*) as a robust *in vivo* regulator of α-Syn levels(10). *Acsbg1* knockdown significantly reduces α-Syn levels in both cultured cells and mouse brains. Furthermore, an independent study has reported increased *Acsbg1* expression with α-Syn toxicity in the α-Syn-E3K model with E35K, E46K, and E61K mutations (13). However, despite its effect on α-Syn, the mechanism of how ACSBG1 regulates α-synuclein pathology remains uncertain.

Acyl-CoA Synthetase Bubblegum Family Member 1 (ACSBG1) is a member of the acyl-CoA synthetase (ACS) gene family whose members encode enzymes that catalyze the conversion of fatty acids into fatty acyl-CoA, enabling their subsequent metabolic utilization into lipid synthesis (14). The expression of *ACSBG1* is confined mainly to the brain, adrenal gland, ovary and testis (15, 16). In the brain, it is exclusively expressed in astrocytes (17, 18). *ACS* gene families are sub-grouped into five categories: short-chain, medium-chain, long-chain, very-long-chain, and the bubblegum family, which includes *ACSBG1* and *ACSBG2* (19). Amongst these, the bubblegum family is known to process long (C16-C20) to very-long-chain fatty acids (C22 or longer) (15, 20). Despite its pivotal role in fatty acid metabolism, the functional contribution of ACSBG1 in brain and to neurodegenerative disease mechanisms remains largely unclear.

Alpha-Synuclein pathology primarily affects neurons, but growing evidence now highlights an increasingly important role of glia—particularly astrocytes, in the progression and worsening of PD pathology. Astrocytes, the most abundant glial cells in the central nervous system, have emerged as pivotal contributors to the progression of neurodegenerative diseases, exerting both protective and pathological effects (21). Astrocytes are dynamic regulators of neuronal function, providing metabolic support to neurons through lactate shuttling, and lipid processing through fatty acid storage and cholesterol transport (22, 23). However, in neurodegenerative conditions such as PD, astrocytes undergo pathologic reactive transformations, shifting from a protective to a detrimental state, contributing to disease progression through mechanisms involving neuroinflammation, oxidative stress, and loss of homeostatic functions (24). Recent genome-wide association studies identified multiple AD and PD risk factor genes, such as *APOE* and *GBA*, that are linked to lipid homeostasis in the cell or between neuron and glial cells (25-27). Also, excessive lipid droplet accumulation in neurons and glia is a marker of dysregulated homeostasis in neurodegenerative diseases (28, 29). Lipid-metabolizing enzymes not only control the balance between lipid storage and utilization, but also directly regulate inflammatory signaling cascades, mediated by bioactive lipids such as lysophospholipids, sphingosine and hexosylceramides that can either protect or damage neighboring neurons (30-32). Targeting these alterations through gene therapy or pharmacological approaches could pave the way for the development of novel treatments for PD.

In this study, we elucidate the mechanism by which ACSBG1 regulates α-Syn and identify ACSBG1 as a previously unrecognized therapeutic target for PD. Mechanistically, we show that ACSBG1 regulates α-Syn levels by modulating inflammation in TNF-α/IL-1α-induced reactive astrocytes. Conditioned medium (CM) from reactive astrocytes elevated total and phosphorylated α-Syn levels in primary neurons, but such an effect was not observed in medium from likewise treated *Acsbg1* knockout (KO) astrocytes. We identified that both chemokines and long chain d20:1 sphingosines were elevated in the reactive WT astrocyte conditioned medium, but loss of *Acsbg1* attenuates this induction. Furthermore, in α-Syn transgenic mice, *Acsbg1* KO reduces astrocyte reactivity, lowers α-Syn pathology, and improves PD-like motor deficits. These findings indicate that ACSBG1 facilitates astrocyte activation and the secretion of pathogenic astrocyte-derived factors to accelerate α-Syn pathology in PD.

## Results

### ACSBG1 deletion suppresses inflammation-induced increase in total and phosphorylated α-Syn by limiting cytokine release from reactive astrocytes

Given the astrocyte-enriched expression of ACSBG1, we proposed that it might regulate α-Syn levels in a non–cell-autonomous manner through modulation of astrocyte–neuron communication. To investigate the role of ACSBG1 in astrocyte-to-neuron paracrine signaling, we employed a conditioned medium transfer paradigm. We first generated reactive astrocytes by treating primary mouse astrocyte cultures with a cytokine cocktail (TNF-α and IL-1α), which drives microglial-induced inflammatory activation observed in PD (33, 34) (Figure 1A). We treated mouse primary neurons with astrocyte conditioned medium and found that conditioned medium from reactive wild-type (WT) astrocytes induced a robust and significant increase in in total α-Syn level and α-Syn phosphorylation at Serine 129 (pS129-α-Syn), which is a major pathological posttranslational modification seen in postmortem human PD tissue (35). In contrast, conditioned medium from *Acsbg1*-knockout (KO) astrocytes did not induce this pathological change (Figure 1B-E). To identify which factors in the conditioned medium induce neuronal α-Syn changes, we activated WT or *Acsbg1* KO astrocytes with pro-inflammatory cytokines (TNF-α/IL-1α) and analyzed both the intracellular response and the resulting secretome (Figure 2A). Transcriptomic analysis via RNA-seq revealed that *Acsbg1* KO significantly modulated the broader TNF-signaling pathway (Figure 2B and Supplementary file 1). This change in intracellular signaling directly impacts the cell’s secretome. Analyzing the extracellular conditioned medium via a large cytokine antibody array shows a marked reduction in the release of pro-inflammatory mediators from cytokine-activated *Acsbg1* KO astrocytes compared to WT controls (Figure 2C, Supplementary Figure 1). The heatmap shows that *Acsbg1* depletion reduced the overall inflammatory response. Several specific factors were identified as being significantly elevated in the WT secretome but strongly reduced in the *Acsbg1* KO: key among them were MIP3α, RANTES, and IL-6, which were selected for further validation. To prove that these identified cytokines cause neuronal changes, we individually supplemented purified RANTES, MIP3α, and IL-6 back into a normal culture medium and applied them to primary neurons. As shown by the Western blot, the presence of any of these three individual cytokines was sufficient to significantly elevate neuronal levels of total α-Syn and, more dramatically, phosphorylated pS129-α-Syn (Figure 2D and E). These findings demonstrate that depletion of *Acsbg1* in reactive astrocytes prevents inflammation-driven phosphorylation of neuronal α-Syn by restricting the release of specific pro-inflammatory cytokines like RANTES, MIP3α, and IL-6.

**Figure 1.**
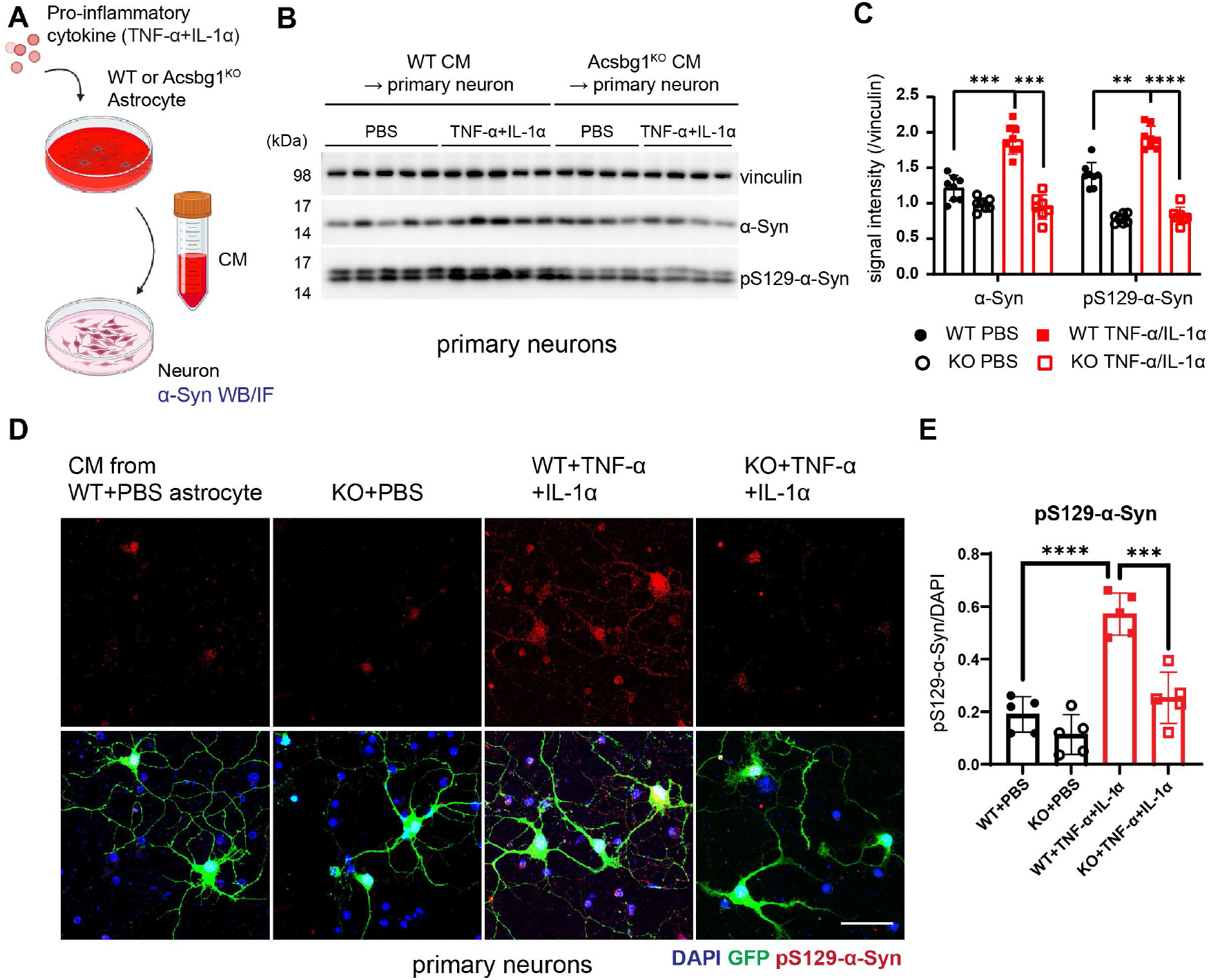
*Acsbg1* deficiency in astrocytes attenuates neuronal *α*-synuclein accumulation and phosphorylation induced by inflammatory conditioned media. (A) Schematic diagram of the astrocyte conditioned media (CM) transfer assay. Wild-type (WT) or Acsbg1 knockout (KO) astrocytes were treated with TNF-α/IL-1α or PBS, and the resulting conditioned medium was applied to primary neurons (B) Representative immunoblot of α-Syn and pS129-α-Syn protein levels in primary neuron lysates following conditioned medium treatment. Vinculin serves as a loading control **(C)** Quantification of α-Syn and pS129-α-Syn band intensities normalized to vinculin. n = 8 per group, **p < 0.01, ***p < 0.001, ****p < 0.0001. (D) Representative immunocytochemical images of pS129-α-Syn (red) in primary neurons treated with conditioned medium from WT+PBS, KO+PBS, WT+TNF-α/IL-1α, or KO+TNF-α/IL-1α astrocytes. GFP (green) marks individual neurons; DAPI (blue) labels nuclei. Scale bar, 50 μm. (E) Quantification of pS129-α-Syn fluorescence intensity normalized to DAPI signal across treatment groups. n = 5-6 per group. ***p < 0.001, ****p < 0.0001.

**Figure 2.**
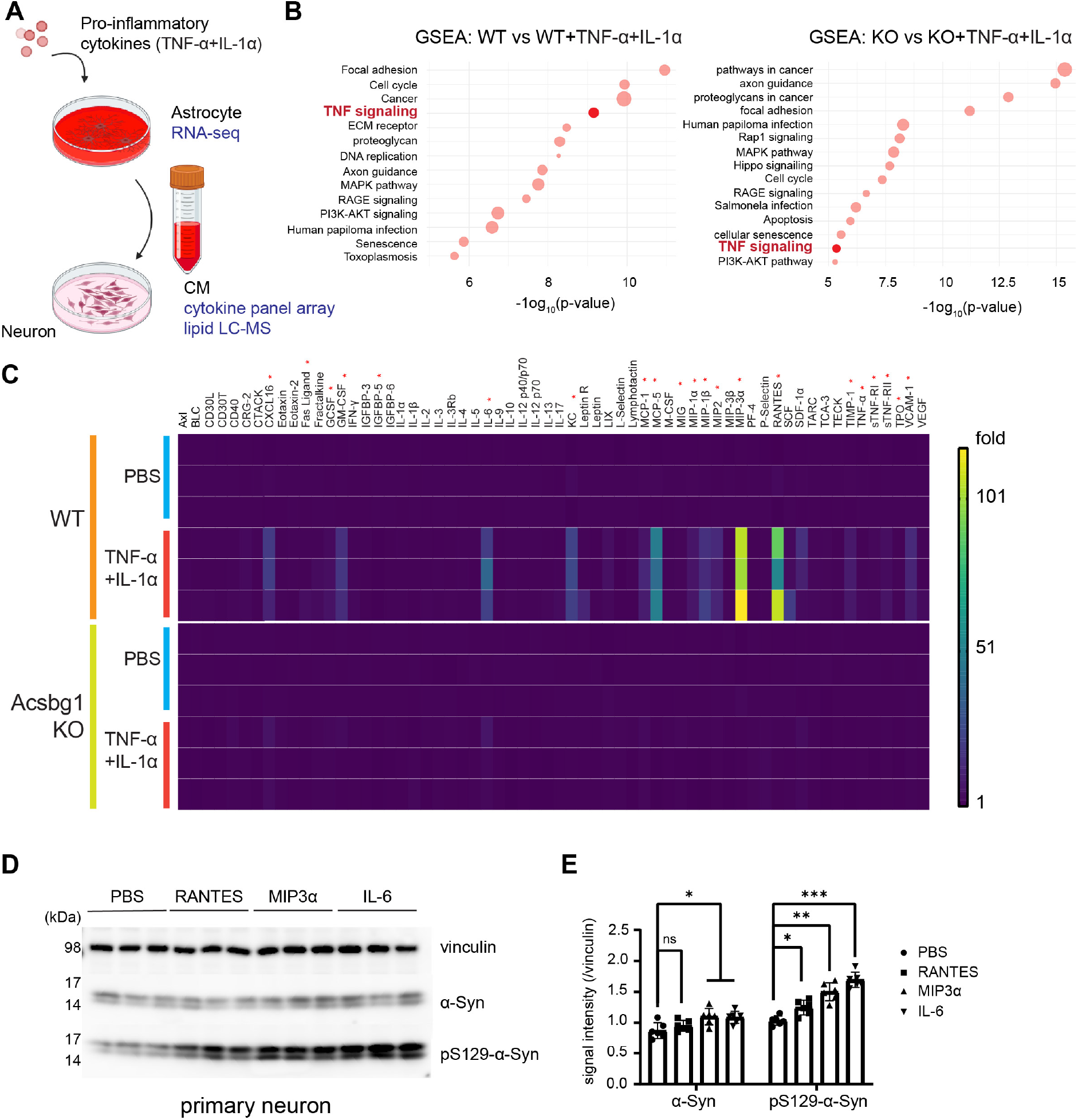
*Acsbg1* deficiency attenuates the inflammatory secretome of TNF-*α/*IL-1*α*-stimulated reactive astrocytes. (**A**) Schematic of the experimental workflow. WT and Acsbg1 KO astrocytes were treated with PBS or TNF-α/IL-1α, followed by RNA sequencing of astrocytes and secretome profiling (cytokine array and lipid LC-MS) of conditioned medium applied to primary neurons. (**B**) Gene Set Enrichment Analysis (GSEA) bubble plots comparing PBS vs. TNF-α/IL-1α-treated WT (left) and Acsbg1 KO (right) astrocytes. TNF signaling (highlighted red) shows a reduced enrichment rank in KO astrocytes relative to WT, indicating attenuated inflammatory pathway activation. (C) Heatmap of cytokine, chemokine, and growth factor levels measured by cytokine array in conditioned medium from WT and Acsbg1 KO astrocytes treated with PBS or TNF-α/IL-1α. Color scale represents fold change relative to the minimum signal (1–101 fold). (D) A representative immunoblot of α-Syn and pS129-α-Syn in primary neuron lysates following treatment with PBS, RANTES, MIP3α, or IL-6 supplement to KO astrocyte conditioned medium. (E) Quantification of α-Syn and pS129-α-Syn band intensities normalized to vinculin. n = 6 per group. *p < 0.05, **p < 0.01, ***p < 0.001; ns, not significant.

### ACSBG1 depletion alters the lipid secretion from astrocytes, including sphingosine, a key regulator of α-Syn

ACSBG1’s known function is to process fatty acids into fatty acyl-CoAs for lipid synthesis(15, 36, 37), accordingly we hypothesized that it might critically shape the composition of astrocytic secretome by modulating intracellular lipid metabolic pathways. To test this, we analyzed the lipid profiles in conditioned medium from WT and *Acsbg1*-deficient astrocytes with or without pro-inflammatory cytokines. While overall lipid classes were similar between reactive astrocytes regardless of *Acsbg1* status (Supplementary file 2), specific long-chain lipids, including sphingosine, LBPA, and hexosylceramide, were significantly upregulated in cytokine-activated astrocytes, compared with control astrocytes. Notably, *Acsbg1* KO restored these lipids to levels comparable to those in control astrocytes (Figure 3A). Building on evidence that ACSBG1 regulates the processing of sphingosine-derived fatty acids, we next sought to identify which specific chain lengths of sphingosine can drive the neuronal α-Syn pathology that is characteristically blocked by *Acsbg1* deficiency. Our lipidomic analysis in Figure 3A indicates that *Acsbg1* KO prevents the pathogenic accumulation of these specific lipids, so we performed an “add-back” experiment. We supplemented conditioned medium from activated *Acsbg1* KO astrocytes (which is low in these specific lipids) with chemically synthesized sphingosine with different chain length ranging from C16:1 to C22:1. We applied this supplemented conditioned medium to primary neurons and monitored the phosphorylation of α-Syn at serine 129 (pS129-α-Syn), a marker of pathological aggregation. Neuronal response, as visualized by immunofluorescence, showed that C16:1 and C18:1 sphingosine did not induce pathology. However, supplementation with C20:1 and C22:1 sphingosine significantly and selectively increased neuronal pS129-α-Syn levels, directly demonstrating their pathogenic potential (Figure 3B and C). Together, these results demonstrate that ACSBG1 is a critical molecular checkpoint in the astrocytic inflammatory response, controlling the production and subsequent secretion of specific long-chain sphingosine species that act as direct mediators to drive neuronal α-synucleinopathies.

**Figure 3.**
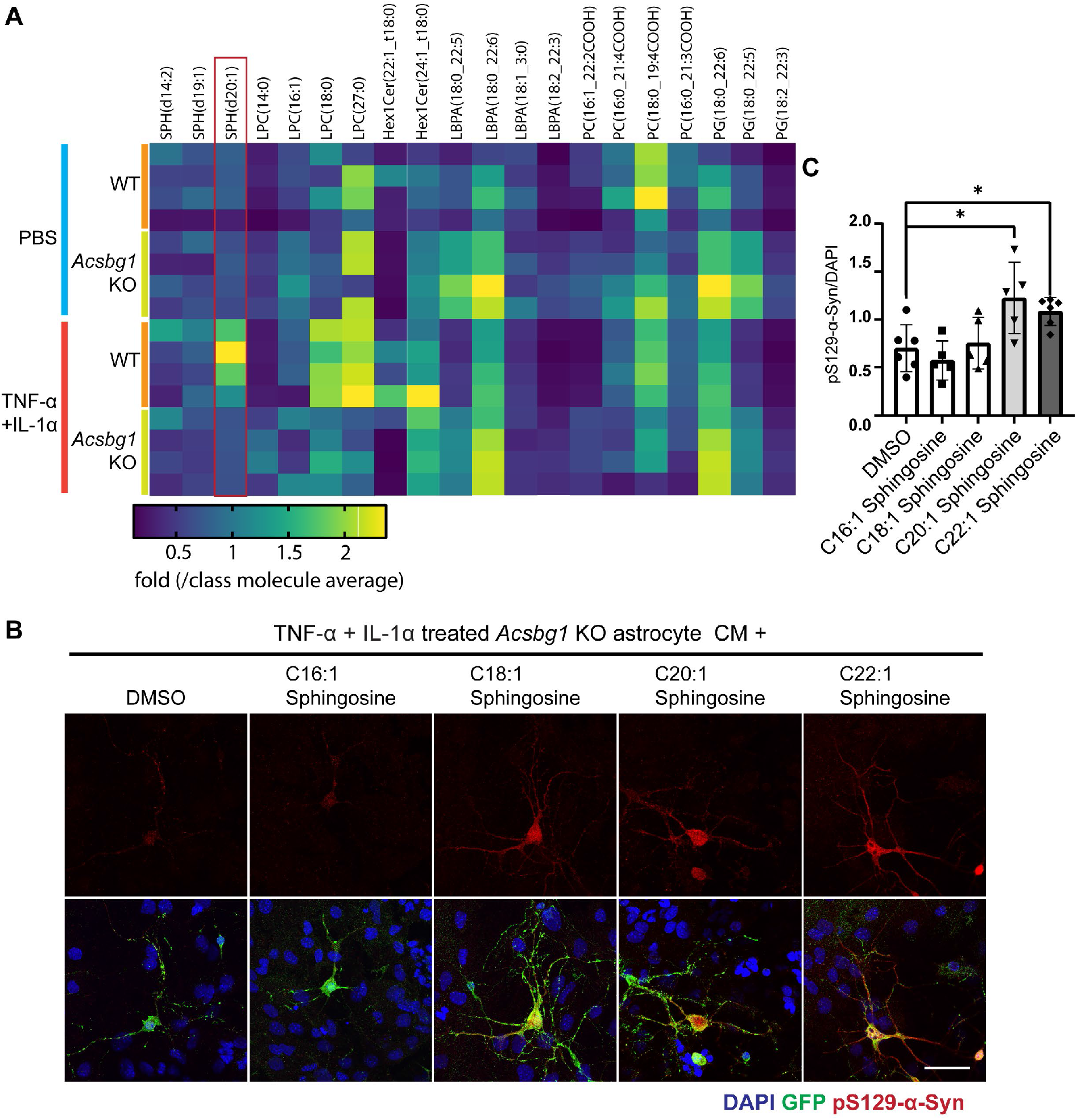
*Acsbg1* KO astrocytes secrete reduced levels of long-chain sphingosines that promote neuronal *α*-synuclein phosphorylation. **(A)** Lipidomics heatmap of lipid species detected in conditioned media from WT and Acsbg1 KO astrocytes treated with PBSor TNF-α/IL-1α. Values represent fold change normalized to the class molecule average. SPH(d20:1) (red box) is notably elevated in TNF-α/IL-1α-treated WT astrocytes and reduced in KO astrocytes. (**B**) Representative immunofluorescence images of pS129-α-Syn (red) in primary neurons treated with Acsbg1 KO conditioned medium supplemented with DMSO (vehicle) or medium- to very long-chain sphingosines (C16:1–C22:1). GFP(green) marks individual neurons; DAPI (blue) labels nuclei. Scale bar, 50 μm. (**C**) Relative quantities of pS129-α-Syn signal intensities compared to DAPI. n = 5–6 per group. *p < 0.05.

### ACSBG1 enzyme activity alters astrocyte status and influences neuronal α-Syn pathology

To determine whether the astrocyte-mediated neuronal α-Syn phosphorylation was dependent on *Acsbg1’s* enzymatic activity, we treated astrocytes with the pan-acyl-CoA synthetase (ACS) inhibitor Triacsin C, followed by stimulation with TNF-α and IL-1α. We observed that Triacsin C-treated reactive astrocytes induced significantly less α-Syn phosphorylation in primary neurons compared to vehicle-treated controls (Figure. 4A-D). To further validate if this effect is exclusively governed by ACSBG1, we generated the catalytically impaired mutant *ACSBG1* (K701A). Biochemical analysis confirmed that ACSBG1 K701A showed a deficit in acyl-CoA synthetase (ACS) activity toward the C18:1 fatty acid substrate (Figure 4E). We then performed a rescue experiment by introducing either WT human *ACSBG1* or the K701A mutant into *Acsbg1* KO astrocytes. Reintroduction of WT ACSBG1 led to similar deleterious effects on neuronal α-Syn pathology. Conditioned medium from *Acsbg1* KO astrocytes reconstituted with WT ACSBG1 could induce robust neuronal α-Syn phosphorylation. In contrast, conditioned medium from *Acsbg1* KO astrocytes expressing an enzymatically inactive ACSBG1 mutant (K701A) failed to induce this effect, resembling *Acsbg1* KO astrocytes (Fig. 4F, G). These data indicate that *Acsbg1* modulates the inflammatory output of reactive astrocytes in an enzymatic activity-dependent manner.

**Figure 4.**
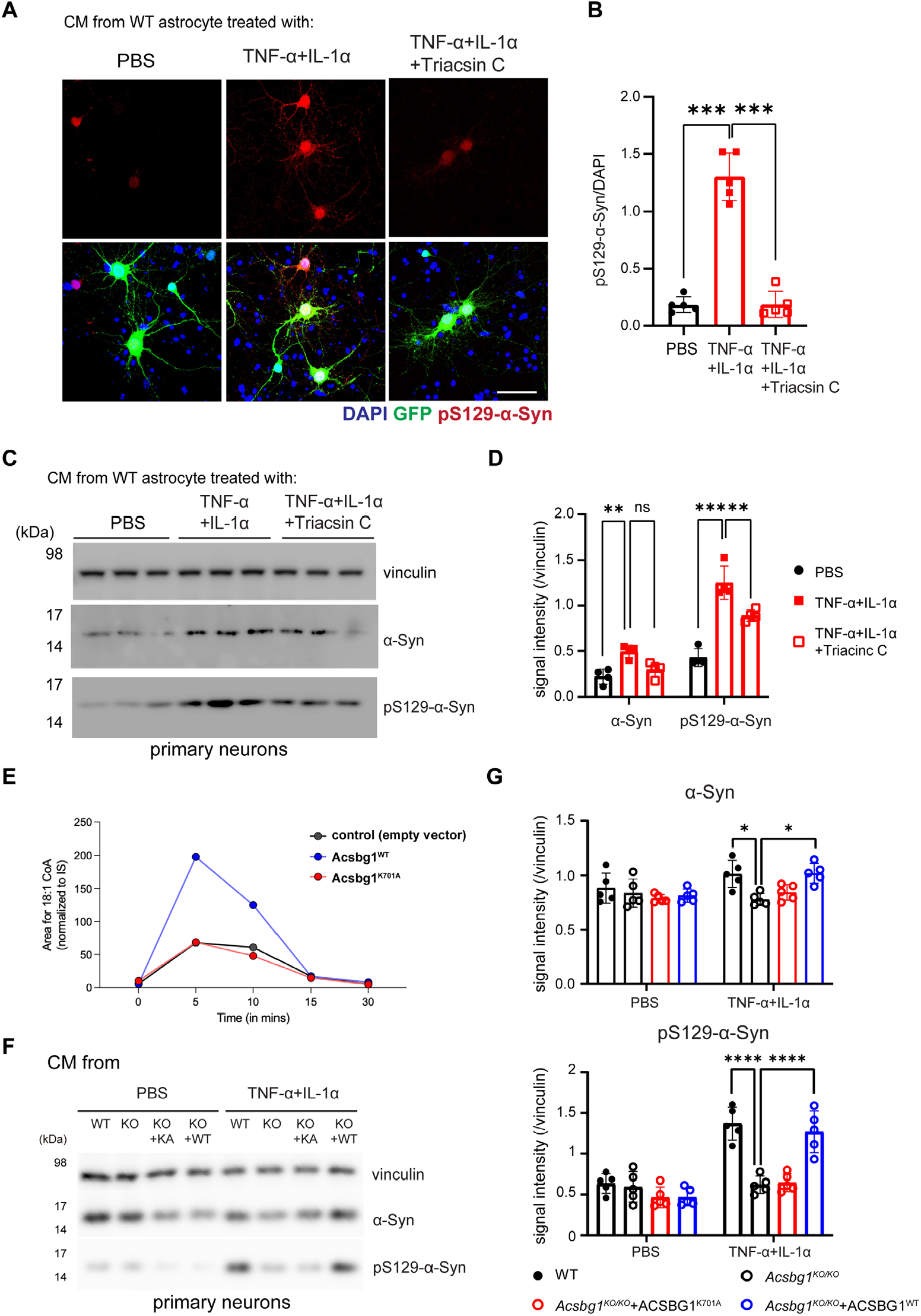
Enzyme activity of Acsbg1 alters the secretion of inflammatory factors from astrocytes. (**A**) Representative images of pS129-α-Syn in neurons treated with conditioned media from cytokine induced astrocytes with or without ACS inhibitor (Triacsin C) (**B**) Relative quantities of pS129-α-Syn signal intensities compared to DAPI. n=5~6 for each sphingosine treatment. n=5, p***<0.001. (**C**) Immunoblot of α-Syn and pS129-α-Syn in neurons treated with conditioned media from cytokine induced astrocytes with or without ACS inhibitor (Triacsin C). (**D**) Quantification of α-Syn and pS129-α-Syn intensities compared to vinculin. n=3, p**<0.01, p***<0.001, ns, not significant. (**E**) Enzyme activity of WT and K701A mutant Acsbg1 protein measured by fatty acyl CoA measurement. (**F**) Immunoblot of α-Syn and pS129-α-Syn in neurons treated with conditioned media from WT or Acsbg1 KO astrocytes expressing Acsbg1 enzyme-dead mutant or wild type Acsbg1. (**G**) Quantification of α-Syn and pS129-α-Syn intensities compared to vinculin. n=5, p*<0.05, p****<0.0001.

### Acsbg1 knockout partially rescues motor impairment in Thy1α-Syn mice

To determine if the inhibitory effect of *Acsbg1* loss on α-Syn accumulation and neuroinflammation observed *in vitro* is preserved in an *in vivo* system, we crossed Acsbg1 KO mice with the Thy1α-Syn (Line 61) transgenic (TG) mouse model. This widely used PD model overexpresses human α-Syn and recapitulates key Parkinson’s disease-associated features, including progressive neuropathology, neuroinflammation, and motor deficits. We performed a comprehensive battery of behavioral tests to assess stereotypic behavior and motor coordination (38). At 10 weeks of age, wild-type (WT) and *Acsbg1*-knockout (KO) control mice displayed normal behavior in the open-field test. In stark contrast, Thy1-α-Syn TG (TG) mice on the wild-type background showed a marked increase in aberrant behavior, with significantly elevated counts for both stereotypic movement (Figure 5A) and vertical rearing activity (Figure 5B). Crucially, the additional deletion of *Acsbg1* in the TG mouse background (Thy1-α-Syn TG;*Acsbg1*^*KO/KO*^, TG;KO) led to normalization of these aberrant behavioral phenotypes. At 10 weeks, the TG;KO double-mutant mice exhibited stereotypic counts that were significantly reduced relative to TG mice alone and indistinguishable from controls. Similarly, the pathological increase in vertical activity was completely absent in the TG;KO mice, which showed a dramatic reduction in this measure back to control levels. Next, we evaluated motor coordination with the pole test and the parallel rod floor test. On the parallel rod floor test, which measures fine motor control by quantifying the number of footfalls per meter, 10-week-old TG mice made significantly more errors compared to wild-type controls (Figure 5C). This deficit was robustly mitigated by *Acsbg1* deletion, with TG;KO mice showing significantly fewer foot slips than TG-only mice. Performance in the pole test, a dexterous coordination task where higher time denotes poorer performance, followed a similar pattern (Figure 5D). TG mice took significantly longer to turn around and descend the pole at 10 weeks compared to WT mice. *Acsbg1* deletion significantly rescued this phenotype, with TG;KO mice exhibiting reduced turn-down times that were faster than those of TG-only mice and closer to control values. Improvements in the pole test were also notable at 5 weeks of age. Despite these clear improvements in specific motor tasks, performance in certain motor tests, such as rotarod and grip strength, was severely impaired in all transgenic mice from the earliest measurable point (5 weeks), making further discrimination difficult (Figure S3A-B). Collectively, these robust *in vivo* data demonstrate that Acsbg1 deletion rescues the pathological elevation of stereotypic behavior and ameliorates specific motor coordination deficits driven by α-synuclein pathology, confirming a pivotal role for Acsbg1 in the progression of Parkinson’s disease-like behavioral deficits in this mouse model.

**Figure 5.**
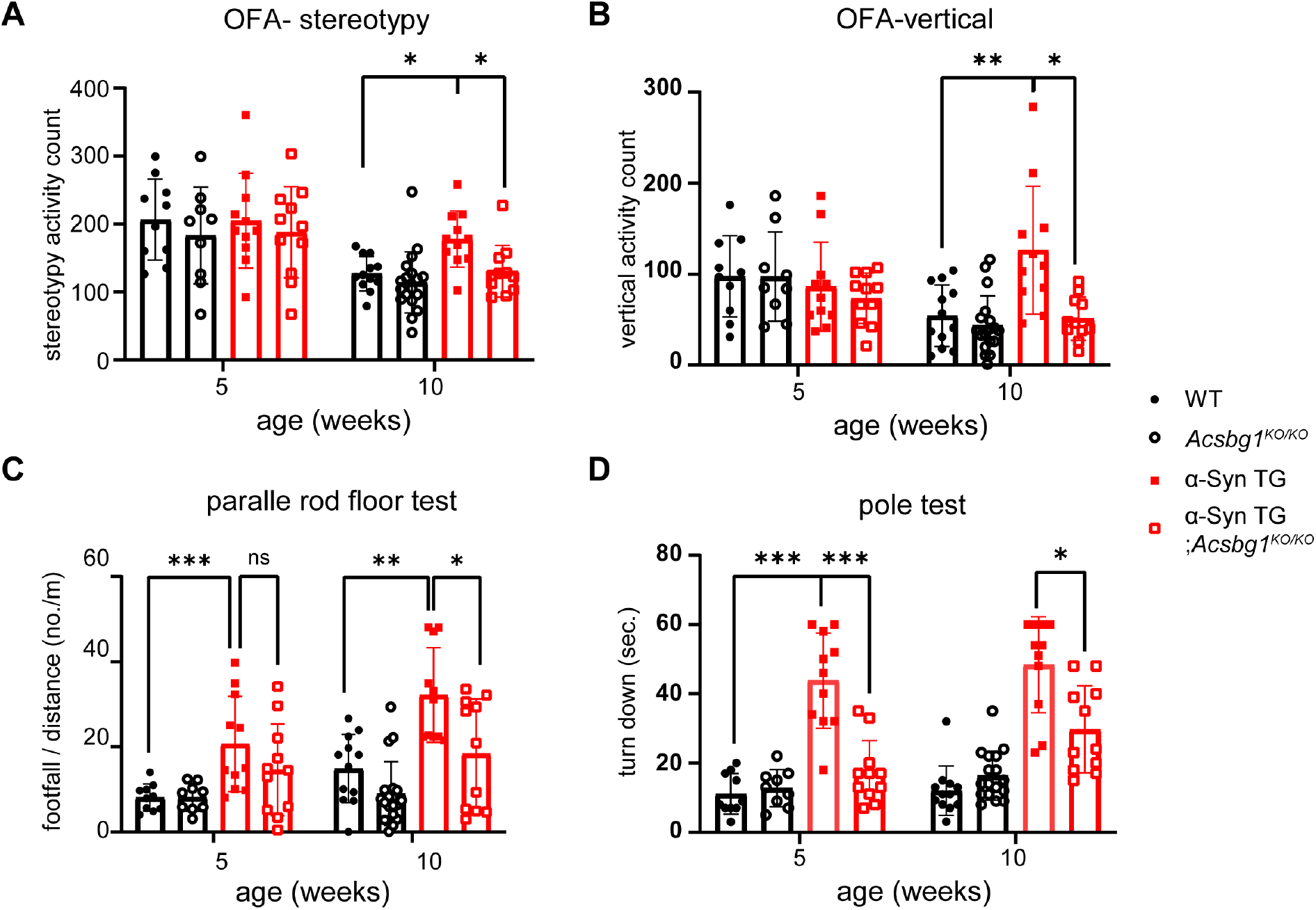
*Acsbg1* KO recovers certainmotorbehaviordeficits shownin *α*-syn transgenic mice. We compared the activity and motor function of WT or Thy1-α-syn TG mouse on the Acsbg1 KO background at 5 and 10 weeks of age. WT (n=10 for 5 weeks, n=12 for 10 weeks), *Acsbg1 KO/KO* (n=9 for 5 weeks, n=18 for 10 weeks), Thy1-α-Syn TG (n=11 for 5 weeks, n=11 for 10 weeks), Thy1-α-Syn TG; Acsbg1 KO/KO (n=11 for 5 weeks, n=11 for 10 weeks). (**A**) Stereotypy in the open field test. (**B**) vertical activity in the open field activity test. (**C-D**) Motor coordination was assessed by parallel rod floor test (**C**) and pole test (**D**). * p<0.05, ** p<0.01, ***p<0.001. All mice used for the behavior test are male mice because Thy1-α-syn TG construct is on the X chromosome.

### Acsbg1 KO reduces inflammation and α-Syn level in vivo

To determine whether the regulatory role of Acsbg1 on α-Syn observed *in vitro* extends to an *in vivo* system, we measured the total and phospho-α-Syn levels at 20 weeks. TG mice showed markedly elevated levels of total and pS129-α-Syn in the cortex (Figure 6A) and midbrain regions including the substantia nigra (Figure 6B), compared with age-matched WT littermates. These levels were significantly reduced on an *Acsbg1* null background (TG;KO) (Figure 6C-F). Furthermore, the TG mice exhibited widespread pathological astrocyte activation, as indicated by robustly elevated expression of the astroglial marker GFAP in both the cortex and substantia nigra compared to WT controls. Strikingly, both this reactive astrogliosis and the associated α-Syn phosphorylation were substantially attenuated in the *Acsbg1*-deficient TG mice (Figure 6G-J, supplementary Figure 3). These findings suggest that mitigating *Acsbg1*-dependent astrocyte reactivity directly reduces the α-Syn pathology. Finally, we examined the downstream impact on synaptic integrity. Thy1-α-Syn TG mice displayed a notable reduction in synaptic density, as evidenced by decreased PSD95 levels in both the cortex (Figures 7A and 7B) and the substantia nigra (Figures 7C and 7D). However, *Acsbg1* deletion prevented this synaptic loss; the TG;KO mice maintained PSD95 expression at levels comparable to WT controls across these regions. Together, these *in vivo* findings demonstrate that *Acsbg1* is a critical mediator of reactive astrogliosis that ultimately drives neuronal α-Syn aggregation and subsequent synaptic degeneration.

**Figure 6.**
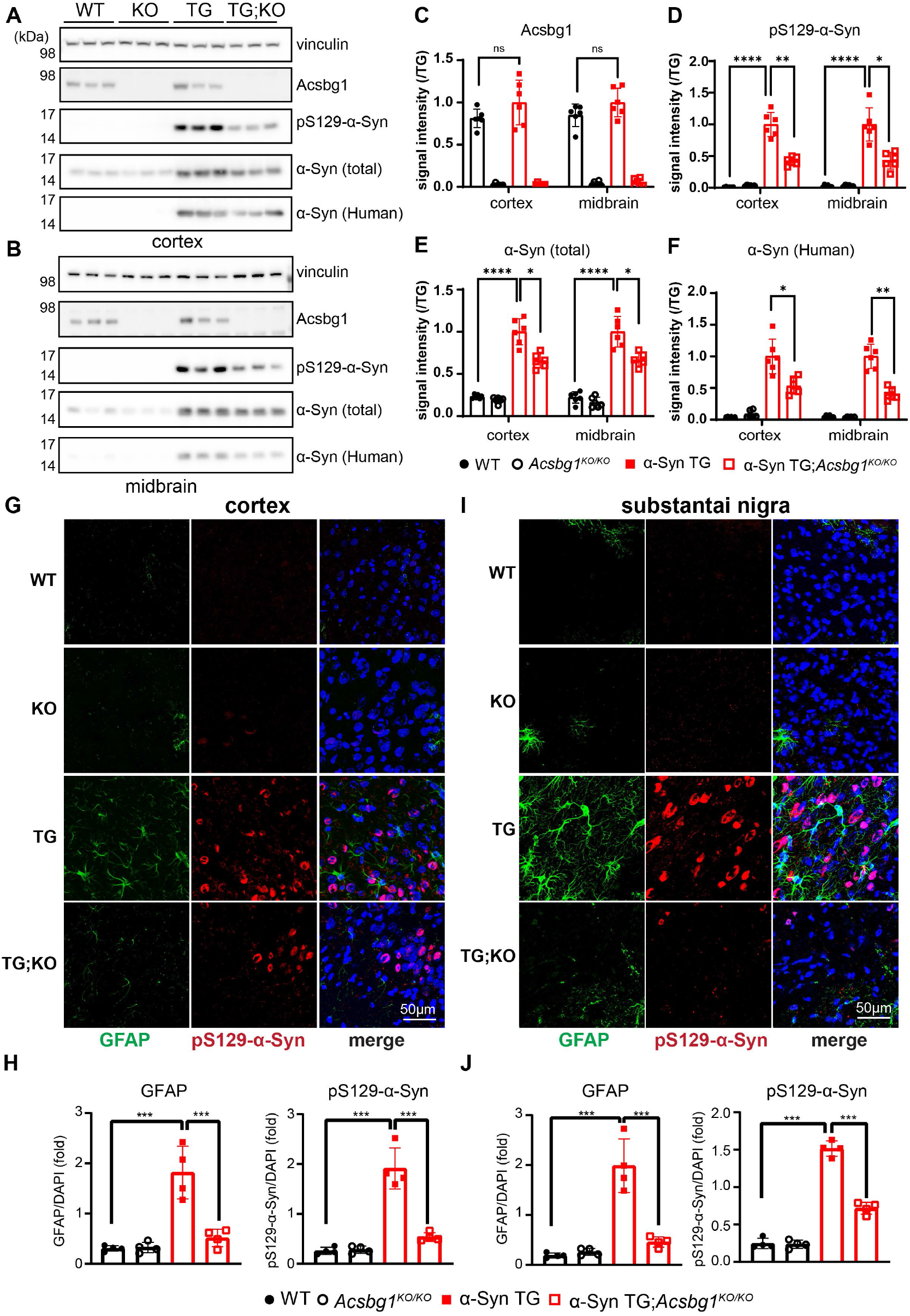
*Acsbg1* KO reduces total and phosphor Ser129 *α*-syn levels with reduced astrocyte activation in *α*-syn transgenic mice. **(A-F)** Immunoblot of Acsbg1 **(C)**, pS129-α-Syn **(D)**, α-Syn **(E)** and human α-Syn **(F)** in cortex **(A)** and midbrain **(B)** from wild type (**WT**), *Acsbg1*^*KO/KO*^ (**KO**), Thy1-α-Syn TG (**TG**), and Thy1-α-Syn TG ; *Acsbg1*^*KO/KO*^ (**TG;KO**) mice. Quantification of protein was normalized with intensities compared to protein level of TG mice. n=6, p*<0.05, p**<0.01, ****p<0.0001, ns, not significant **(G, I)** Representative images of astrocyte activation in the cortex (**G**) and substantia nigra (**I**) visualized with pSer129-α-Syn and GFAP double immunostaining. (**H, J**) Relative quantities of GFAPin cortex(**H**) and substantia nigra(**J**). GFAPsignal intensities compared to DAPI. n=4. p***<0.005.

**Figure 7.**
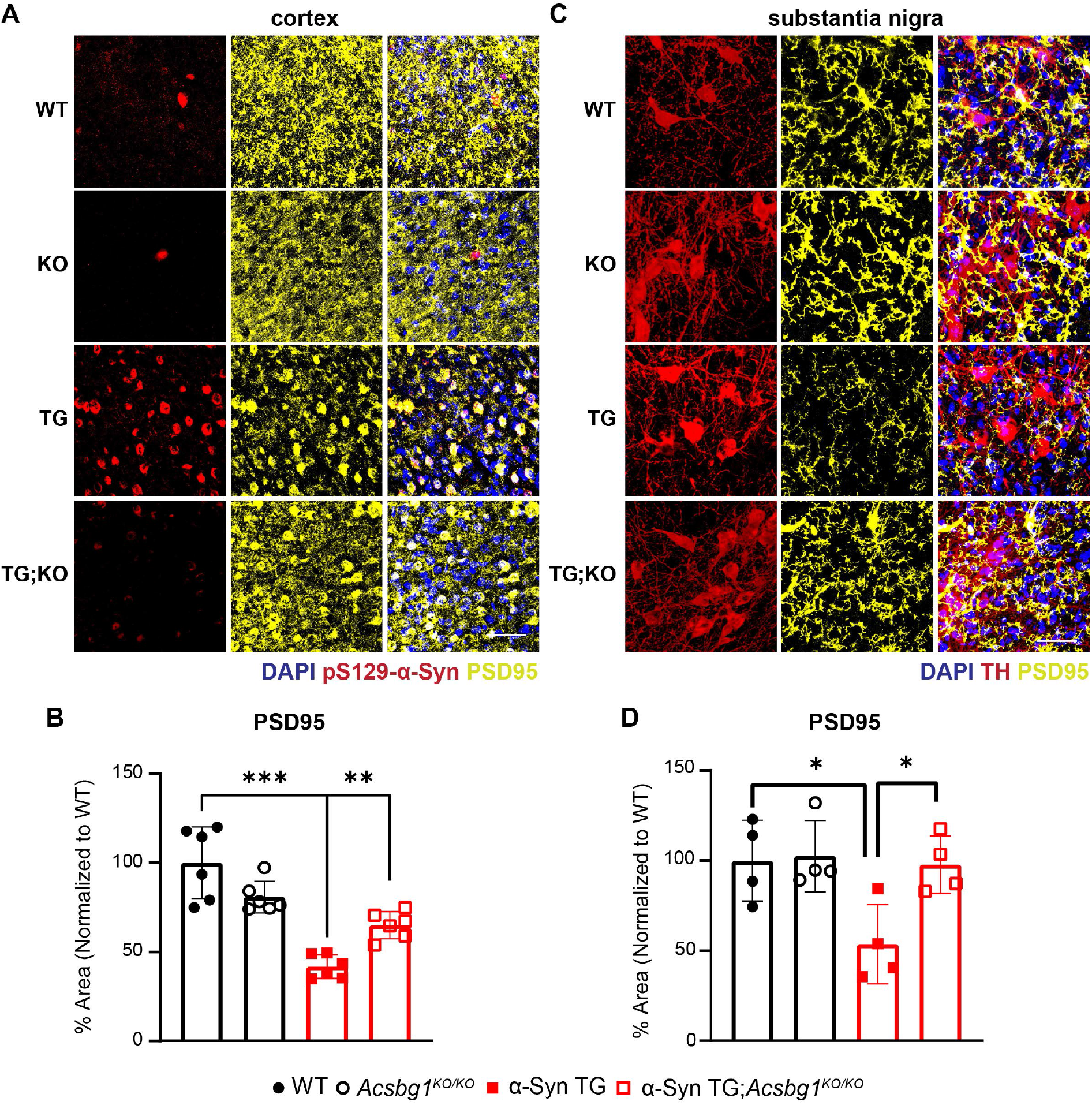
*Acsbg1* KO rescues reduced synaptic density in *α*-Syn transgenic mice. Post synaptic density is assessed in 20-weeks-old WT, Acsbg1 KO/KO, Thy1-α-Syn TG, and Thy1-α-Syn TG ; Acsbg1 KO/KO mice (n=4-6). Scale bar (white): 50µm. (**A**) Representative images of synaptic density in the cortex visualized with pSer129-α-Syn and PSD95 double immunostaining. (**B**) Relative quantities of PSD95 signal intensities compared to DAPI. p**<0.01, p***<0.001. (**C**) Representative images of synaptic density in the substantia nigra visualized with TH and PSD95 double immunostaining. (**D**) Relative quantities of PSD95 signal intensities at substantia nigra compared to DAPI. p*<0.05.

## Discussion

While the importance of astrocyte-neuronal crosstalk in central nervous system homeostasis is well established, the specific glial factors that corrupt this interaction during neurodegeneration remain elusive. In this study, we identify the astrocyte-enriched lipid metabolic enzyme ACSBG1 as a central mediator of α-Syn pathogenesis. Mechanistically, our findings indicate that ACSBG1 functions as a critical metabolic switch; by coupling lipid processing to the inflammatory secretome, ACSBG1 activity actively drives astrocytes into a neurotoxic, reactive state. Importantly, we demonstrate that targeted *Acsbg1* deletion uncouples this deleterious crosstalk *in vivo*, significantly reducing both total and pathological phosphorylated α-Syn accumulation, resolving reactive astrogliosis, and rescuing motor coordination deficits in Thy1-α-Syn transgenic mice. Ultimately, these data position ACSBG1 as a pivotal regulator of Parkinsonian α-synucleinopathies, linking astrocytic lipid metabolism directly to neuronal vulnerability.

Recent genome-wide association studies (GWAS) have heavily implicated lipid metabolism networks in neurodegenerative disease pathogenesis (29, 39-42). While much of the initial evidence comes from Alzheimer’s disease (AD), in which genes such as *APOE, CLU* (ApoJ), *PICALM, TREM2*, and *ABCA7* promote disease-associated glial activation along with lipid droplet (LD) accumulation and lipid dysregulation across both neurons and glia (43-48), similar metabolic abnormalities are increasingly recognized in PD as well. For instance, enzymes that regulate saturated fatty acids, such as SCD1 and SCD5, significantly alter α-Syn aggregation in both yeast and mammalian neuronal models (49, 50). Furthermore, *GBA1* mutations as the most significant genetic risk factor for PD cause the accumulation of glucosylceramide (GlcCer) and glucosylsphingosine (GlcSph) in neurons (51), and simultaneously drive severe dysregulation of lipid homeostasis in astrocytes (52). SPTSSB, a gene that is key regulator of de novo sphingolipid biosynthesis, has been recently identified as risk factor of PD (53). Consistent with this metabolic stress response, previous studies have shown that ACSBG1 expression is upregulated in oxidatively stressed or lipid-droplet-forming astrocytes (23). However, the specific regulatory networks within astrocytes that control astrocyte reactivity and secretion of disease-modifying lipids and cytokines remain elusive. Our findings position Acsbg1 as a central metabolic regulator linking astrocytic lipid metabolism to α-Syn pathology.

Reactive astrocytes secrete pro-inflammatory cytokines and inflammatory mediators, including RANTES, MIP-3α and IL-6, which are elevated in PD patient plasma and robustly induce α-Syn phosphorylation (54-57). Our data show that exogenously applying these specific cytokines directly increases neuronal α-Syn phosphorylation (Figure 2), implicating their role in astrocyte-neuron crosstalk. Beyond inflammatory mediators, we also identified a novel lipid-mediated mechanism that exacerbates neuronal α-Syn pathology. We found that specific sphingolipid species, notably d20:1 sphingosine, were significantly upregulated in TNF-α/IL-1α-induced reactive wild-type primary astrocytes conditioned medium but normalized upon depletion of *Acsbg1* (Figure 4). Furthermore, extracellular application of these long-chain sphingosine species to conditioned medium from Acsbg1-deficient reactive astrocytes was sufficient to induce neuronal pS129-α-Syn (Figure 4). Together, these findings demonstrate that reactive astrocyte-mediated α-Syn phosphorylation occurs through cytokines and lipid metabolites.

In PD, these sphingolipid species provide a critical mechanistic link between glia and neurons. Sphingosine kinases (SphK) convert Sphingosine to sphingosine-1-phosphate (S1P), which acts as a potent signaling molecule. Alternatively, sphingosine is converted to ceramides, which ultimately gives rise to GlcSph and GlcCer (58). Elevated long chain sphingosine including d20:1, have been shown to drive neurodegeneration in mice (30). Although the S1P pathway is normally involved in regulating autophagy and cellular survival (59), disease states fundamentally alter this balance; S1P and SphK levels are highly upregulated in the PD patient’s CSF and have been identified as microglial activators in α-Syn mouse and cultured microglia models (60, 61). Neurotoxic lipid species like GlcSph and GlCer are known to accumulate in lysosomes (62), and both sphingosine and GlcCer actively increase α-Syn oligomerization by enhancing its seeding capacity (63). Prior evidence suggests that ACSBG1 plays a role in sphingosine metabolism (64), directly connecting sphingolipid dysregulation in PD to this gene.

Because ACSBG1 functions as an acyl-CoA synthetase specifically for long- to very-long-chain fatty acids, which serve as essential precursors for complex sphingolipid assembly. Our findings, therefore, identify *Acsbg1* as a critical upstream regulator of the pathogenic lipid cascade. In this study, suppressing ACSBG1 activity prevents the production and secretion of these longer chain neurotoxic sphingolipids. Similarly, ACSL1, another member of the ACS family, which converts long chain fatty acids to fatty acyl-CoA, has also been shown recently to be a critical mediator of microglia reactivity (45). Collectively, these findings highlight a broader role of ACS family genes in coupling glial activation to lipid processing.

We also corroborated prior findings that longer-chain lipids drive α-synucleinopathies. Very long-chain (C22) GlcSph and GlcCer have previously been established as hallmarks of disease (65). As the blood brain barrier can restrict the permeability of longer chain fatty acids and their derivatives due to their hydrophobicity (66), our lipidomic data prioritizes long chain sphingolipid metabolites are derived from astrocytes, and potential modulators of neuronal α-syn pathology. Consistent with prior evidence that demonstrated that very-long-chain sphingolipids enhance synuclein aggregation (65), our data further implicate the role of longer-chain sphingosine species secreted by reactive astrocytes as a potential driver of neuronal α-Syn phosphorylation and accumulation.

The unique biological profile of ACSBG1 makes it a promising therapeutic target. ACSBG1 is predominantly expressed in astrocytes within the central nervous system (17), and its systemic expression is highly restricted (limited primarily to the testis and ovary in humans). Most importantly, *Acsbg1* knockout mice exhibits no overt baseline physiological or behavioral phenotypes. This suggests that the brain can tolerate the loss of *Acsbg1* under healthy conditions, and that pharmacological inhibition of this enzyme could selectively mitigate disease-associated astrocytic reactivity and α-Syn pathology with a wide therapeutic index and minimal off-target toxicity.

While our findings highlight the therapeutic potential of targeting ACSBG1, the precise downstream mechanisms linking its activity to diminished astrocytic reactivity require further investigation. It remains to be determined whether mitigating this enzyme’s activity reduces the intracellular lipid burden within the astrocyte (thereby restoring its resting state), or simply removes the primary sphingolipid stressor, allowing neurons to recover. Additionally, as this study primarily utilized a Thy1-α-Syn model that relies on robust transgenic overexpression, that makes it hard to see the difference in grip strength or rotarod test, which already shows severe deficits from a very early age (5 weeks) (supplementary Figure 3), future studies must evaluate *Acsbg1* deletion in more physiologically relevant models. Testing this mechanism in an α-Syn knock-in mouse model, such as the *Snca*^G51D/G51D^ model, which accurately captures the prodromal symptoms of Parkinson’s disease before severe motor deficits appear, would be ideal (67). It will provide a better interpretation of how targeting astrocytic *Acsbg1* can mitigate (or attenuate) the synucleinopathies during the progression of pathology.

Our findings identify Acsbg1 as a master regulator of astrocyte reactivity and neuronal α-Syn pathology through astrocyte to neuronal signal transfer. We demonstrate that ACSBG1 drives α-Syn phosphorylation and aggregation through the release of inflammatory factors (RANTES, MIP3α, and IL-6), with lipid-mediated secretion of d20:1 sphingosine. Consequently, Acsbg1 emerges as a novel therapeutic target for modulating astrocyte-neuron communication in PD.

## Material and Methods

### Sex as a biological variable

Sex was not considered as a biological variable for the *in vitro* studies. For the mouse experiment, as the α-syn transgene is on the X chromosome, only male offspring were used in the experiments because of females’ variability linked to X-inactivation.

### Animal studies

All mice were housed in a level 3, American Association for Laboratory Animal Science (AALAS)–certified facility on a 14-hour light cycle. Husbandry, housing, euthanasia, and experimental guidelines were approved by the Institutional Animal Care & Use Committee (IACUC) at Baylor College of Medicine.

### Mouse breeding

Acsbg1 KO mice (37) and the Thy1-α-Syn (“Line 61”) (38) mice on C57B6/J background were used for experiments. In experiments using the Thy1-α-Syn, transgenic females on WT or Acsbg1 KO background were bred with wild-type males or with Acsbg1 KO mice.

### Behavioral Assays

We tested 5- and 10-week-old WT, Acsbg1 KO, Thy1-α-Syn TG, and Thy1-α-Syn TG on Acsbg1 KO background mice with their wild-type littermates. A series of behavioral tests were performed as a part of this study. Mice were acclimated to the test environments for 30 minutes before testing, with at least a 30-minute interval between each trial.

### Open-field test

The open field arena is a 40 × 40 × 40 cm (width × length × height) cubical enclosure. The central 20 × 20 cm region of the box was marked by ANY-maze software (Stoelting Co.). The mouse was placed in a corner of the box and recorded by an overhead video camera for 10 minutes. Several measures were analyzed, including total distance travel, mean motor speed, and center/total ratio.

### Pole test

The mice were placed head up near the top of a vertical pole (threaded rod, 50 cm long, 1 cm in diameter), and the test lasted 60 seconds. We counted the number of times the mouse turned its head and whole body downward (turn) and climbed down to the ground. The mice underwent 3 trials and the minimum durations of ‘turn’ and ‘down’ time were measured and used for analysis.

### Grip Strength

A mouse was picked up from the base of a tail and gently lowered towards the net until it grasped the bar of the grip strength meter (Columbus Instruments 0167-8001). The mouse was then gently pulled backward until it released its grip. The maximum pull force at the time the animal released the grip was recorded on a horizontally mounted scale equipped with a drag pointer. Mice underwent three trials and the maximum score across the three trials was used for the analysis.

### Parallel rod floor test

Animals were placed individually into the center of a wire grid laid within an open-field chamber (AccuScan) for 10 minutes. The number of paw slips through the wire grid were recorded and analyzed using ANY-maze (Stoelting). The number of foot slips was normalized to the total distance traveled and then compared across groups.

### Rotarod

Mice were placed on an accelerating rotarod apparatus (Type 7650, Ugo Basile) and allowed to move freely as the cylinder increased from 5 r.p.m. to 40 r.p.m. over a 5-min period. Each trial lasted a maximum of 10 minutes, and the latency to fall was measured when the mouse fell off the rod or rode the cylinder for two consecutive revolutions without regaining control. The maximum time cut-off of the test was 600 seconds. Each training day consisted of four attempts with a 30-minute rest between each trial for 4 days.

### Antibody, compound and construct used for this study

All the antibodies, reagents, animal models, DNA constructs are on the supplementary table 1.

### Protein extraction

Mice were sacrificed by isoflurane inhalation at 20 weeks of age. The cortex, striatum, and midbrain region containing the substantia nigra were harvested and immediately frozen on dry ice. Samples were mixed with 5-10 volumes of modified RIPA buffer (50 mM Tris-Cl, 150 mM NaCl, 1% NP-40, 0.5% sodium deoxycholate, 0.1% SDS) supplemented with 0.5% triton X-100, 1X protease inhibitor, and 1X phosphatase inhibitor buffer, and lysed with a sonication step (20 pulses, output 2.5, duty cycle 30%, 2 sec and 2 sec rest intervals, 5 times). Samples were centrifuged at 20,000 g for 20 min, and the supernatant was collected for use.

### SDS-PAGE and Western blot

Protein samples were loaded on either 10- or 15-well NuPAGE 4-12% Bis-Tris gels. Gels were run in 1X MES/SDS protein running buffer and transferred onto nitrocellulose membranes in Tris-Glycine buffer (25 mM Tris, 190 mM Glycine) at 0.3 amps for 1.5 hours. After being transferred, membranes were blocked in 5% milk in TBS-T for 1 hour and probed with one of the following primary antibodies overnight: mouse anti-vinculin (1:10000), rabbit Acsbg1 (1:3000), mouse anti-α-Syn (1:3000), rabbit anti-pS129-α-Syn (1:3000). Membranes were subsequently washed 3 times in TBS-T for 10 minutes and mouse or rabbit HRP-conjugated secondary antibodies were applied in 5% skim milk in TBST. Following the wash, chemiluminescence was induced using ECL (GE Healthcare, RPN2236) and imaged with an Amersham imager 680 (GE Healthcare).

### Tissue preparation

For immunofluorescence and immunohistochemical experiments, mice were transcranial perfused with PBS followed by 4% paraformaldehyde (PFA). Brains were dissected and fixed in 4% PFA for 2 days, dehydrated for 24 hours in 15% sucrose (w/v, in PBS), followed by a 2-day incubation in 30% sucrose solution (in PBS), all at 4 °C. The brains were then frozen on dry ice in OCT compound (VWR, 25608-930) and sectioned on a cryostat (Leica CM 3050S). Sections were collected at 40 µm thickness and kept in 1X PBS with 0.01% NaN_3_ until ready for use.

### Primary Astrocyte culture

Primary astrocyte culture method was adopted from Schildge et al.(68) with some modifications. Briefly, P3 WT or Acsbg1^KO^ pups were taken and their heads and necks were thoroughly sprayed with 70% ethanol. The brain was then removed and placed in prechilled HBSS solution under a stereomicroscope. Careful dissections were performed to remove cortices from other brain regions, as well as meninges and thereafter transferred to a centrifuge tube containing ice-cold HBSS. Subsequently, trypsin was added to the cortices, and the tubes were placed in a water bath at 37°C for 10 min. Then, the cortical tissue pieces were centrifuged, the supernatant was removed, and astrocyte plating media (DMEM, high glucose + 10% fetal bovine serum + 1% Penicillin/Streptomycin) was added to the tube. A single cell suspension was then prepared, and cells were plated in a poly-D-lysine-coated T75 flask. The media was changed after 2 days of plating and every 3 days thereafter to get an astrocyte-enriched culture. The flask was shaken after a week to remove other non-neuronal cells, including microglia and oligodendrocytes, as described earlier. The cells were then washed, and plating media was added again to the cells.

### Collection of Astrocyte-conditioned media

Astrocyte conditioned medium (CM) was collected from cytokine-treated (TNF-α and IL-1α) and control (PBS) astrocytes. Firstly, astrocytes were treated for 24 h in a serum-free medium with the cytokine cocktail containing TNF-α (50ng/ml) and IL-1α (15ng/ml). Thereafter, the cells were washed with PBS, and fresh complete medium was added to the cells. Subsequently, after 24 h, the media as well as astrocytes were collected and separately stored at - 80°C until further use.

### Primary Neuronal Culture

Primary neuronal culture was done from P0 pups of WT mice using the papain dissociation system (Worthington LK003153). The heads and necks of mice were sprayed with 70% ethanol. Thereafter, brains were dissected under a stereomicroscope, and cortices were separated after removing the meninges. Following cortical resection, tissue from each pup was placed into 0.5 ml of complete neurobasal media (Neurobasal media, 2% B27 supplement, 0.5% Penicillin/Streptomycin and 0.5% GlutaMax) placed in ice and pre-equilibrated with 95% O_2_:5% CO_2_ in a CO_2_ incubator. The complete neurobasal media was removed and cortices were then digested using pre-equilibrated Papain solution for 45 min at 37°C in a thermomixer. The solution was then removed, and pre-equilibrated inhibitor was added to the tissue and incubated in a thermomixer for 5 min at 37°C. Thereafter, the inhibitor was removed and 1 ml of complete neurobasal media was added to each centrifuge tube, followed by trituration (~25 times) using a pipette to make a single cell suspension. Subsequently, the cells were passed through a 40 µm cell strainer and then counted using trypan blue solution in a hemocytometer. Finally, 1 × 10^5^ cells/well were seeded in poly-d-lysine-coated 24-well plates for further experimentation. For immunofluorescence experiments, cells were seeded onto precoated cover glasses placed in 24-well plates. Neurons were allowed to attach to the wells or cover glasses, and the media was removed after 2 h to remove unattached glial cells. Media was changed every 4 days.

### Cytokine and sphingosine treatment of primary neurons

Primary wild-type neurons were cultured for 10 days *in vitro*, after which they were treated for 24 hours with either 20 ng/µL of RANTES, MIP3α, and IL-6, or with astrocyte-conditioned medium derived from TNF-α/IL-1α-stimulated *Acsbg1* KO cells, supplemented with d16:1 to d22:1 sphingosine. Following treatment, neurons were harvested for immunofluorescence and Western blot analysis.*Immunofluorescence*. Immunofluorescence was performed on precoated coverslips in 24-well plates or Floating sections. Briefly, cultured neurons and floating sections of mouse brain were washed thrice in 1X PBS and then incubated with one or two of the following antibodies overnight: mouse anti-α-Syn (1:1000), rabbit anti-pSer129-α-Syn (1:1000), rabbit anti-Iba1 (1:1000), goat anti-GFAP (1:1000), and rabbit anti-Tyrosine Hydroxylase (TH) (1:1000), rabbit anti-post synaptic density 95 (PSD-95) (1:1000).. The next day, sections were washed three times with 1X PBS and stained with secondary antibodies with a designated fluorophore (488 or 555nm) at 1:1000 dilutions *at* room temperature for 3 hours. After three washings, the coverslips were taken and mounted with VECTASHIELD® HardSet™ Antifade Mounting Medium (Vector Laboratories, H-1400-10). Imaging was performed on a confocal microscope (Leica STED TCS SP8X), using LAS X software (Leica) after selecting optimal settings for image capture. All immunofluorescence images presented in this study were taken with the 63X objective lens.

### Inflammation Array

Levels of cytokines, chemokines, growth factors, and other inflammatory mediators were measured using the Mouse Cytokine Antibody Array-Membrane kit (Abcam, MA, USA). Briefly, the array membranes were blocked using a blocking buffer at room temperature for 30 min followed by incubation with conditioned media overnight. Subsequently, membranes were washed with wash buffer I and II provided with the kit and further incubated with biotin-conjugated antibody cocktail at room temperature for 2 h. The membranes were washed again, and HRP-conjugated streptavidin antibody was then added to them for 2 h at room temperature. Finally, the membranes were washed and signal intensity was quantified using chemiluminescence detection buffer, followed by analysis using ImageJ software.

### RNA Extraction and Bulk RNA Sequencing

Astrocytes frozen following the collection of conditioned media were scraped from the surface of flasks and subsequently RNA was extracted from them using the RNeasy Mini Kit (Qiagen #75106), according to the manufacturer’s instructions. Subsequently, RNA concentrations were quantified with the help of a NanoDrop, and the RNA was sent for library preparation and sequencing on the GeneWiz Illumina HiSeq Platform. Raw reads were quality-assessed with FastQC v0.11.9 and trimmed using fastp v0.20.0. Trimmed reads were aligned to the mouse reference genome GRCm38 (GENCODE release M23) using STAR v2.7.11a with a 2-pass alignment strategy. Gene-level counts were obtained using STAR’s --quantMode GeneCounts parameter. Differential expression analysis was performed with DESeq2. Genes with mean counts below 50 across all samples were excluded prior to testing. Differentially expressed genes (DEGs) were defined by an adjusted p-value ≤ 0.05 and |log_2_ fold change| ≥ 0.263 (69). Functional enrichment analysis of significant DEGs was performed using gProfiler to identify enriched KEGG pathways.

### Lipid analysis

All samples were analysed on a Vanquish UHPLC (Thermo Scientific) coupled to a Orbitrap Exploris 240 mass spectrometer (Thermo Scientific #BRE725535). For routine lipid analysis, extracts were separated on a C18 reverse phase column (Pheneomenex, 5 μm, 50 x 4.6 mm #00B-4435-E0) as described(70). Briefly, for positive mode: solvent A= 95:5 (v/v) H2O/MeOH + 0.1% formic acid + 10 mM ammonium formate, solvent B= 60:35:5 (v/v/v) isopropanol (IPA)/MeOH/H2O + 0.1% (v/v) formic acid + 10 mM ammonium formate. For negative mode, solvent A= 95:5 (v/v) H2O/MeOH + 0.1% (v/v) NH4OH, solvent B= 60:35:5 (v/v) IPA/MeOH/H2O + 0.1% (v/v) NH4OH. LC gradient was 60 min, with stable flow rate of 0.5 mL/min starting 5% B for 5 min, linear gradient of B from 5–100% over 50 min, 100% B from min 50–55, and re-equilibration with 5% B for 5 min. The column temperature was maintained at 55 °C. MS parameters were as follows: - heated ESI mode of ionization, sheath gas = 40 units, aux gas = 8 units, sweep gas =1 units, spray voltage = static, positive ion =3,400 V, negative ion = 2500 V, ion transfer tube temp= 320 °C, vaporizer temp=275 °C. Orbitrap resolution at MS1 was set to 120000, scan range 250–1800, RF lens = 60%, AGC target = standard, Orbitrap resolution at MS2 was set to 30000. EASY-IC™ was used for internal calibration. Lipids were searched and aligned for annotations in LipidSearch 5.1 (Thermo Scientific #OPTON-30880) with precursor mass error tolerance of 5 ppm and product mass error tolerance of 10 ppm, retention time deviation tolerance of 0.5 min.

### Generation of Acsbg1^K701A^ mutants

Acsbg1 WT plasmid was procured from GenScript. A potential mutation that interferes with the catalytic activity of the enzyme is predicted by Uniprot.org (https://www.uniprot.org/uniprotkb/Q96GR2/entry); specifically, K701A was predicted to reduce enzyme activity. We mutated K701 to A using Q5-site directed mutagenesis kit (New England Biolabs #E0554S).

### Statistical analysis

For all experiments, comparisons of two groups were performed using Student’s t-test and unequal variance t-test when the group sizes were very different (Figure 6). Comparisons of three or more groups were performed using one-way ANOVA followed by Dunnett’s multiple comparison test. All analyses were conducted using Prism 10 software (GraphPad). Data represents means ± s.e.m.;

### Image creation with Softwares

Figure 1A, Figure 2A and graphical abstract is generated on Biorender. All the image quantification is done with ImageJ (v2.16.0), and all the graphical works are done in Adobe illustrator (v30.4). We have a license to use these programs.

## Data availability

The RNA-seq raw data generated in this study for WT or Acsbg1 KO primary astrocytes have been deposited in the NCBI GEO under accession code GSE332590. All other data supporting the findings of this study are available within the paper and its Supplementary Information.

## Authors’ contributions

Y.K. and H.Z. conceived the project, Y.K, B.V, S.S., H.K.Y., and H.Z. designed the study/experiments,

A.K.V. and H.K.Y. analyzed RNA-seq data, F.J.S. synthesis the lipids used in this research, J.K and S.B. made the DNA constructs for the research; and Y.K wrote the paper and H.Z. edited it.

## Acknowledgements

We appreciate Hamin Lee and Mason Tate for critical comments on the manuscript. We thank the Microscopy Core and Animal Behavior Core supported by the Jan and Dan Duncan Neurological Research Institute (NRI) at Texas Children’s Hospital, and Baylor College of Medicine (BCM)

