## Supplement table 1 and Supplement figure 1-3 for "Astrocytic ACSBG1 depletion improves lipid-cytokine signaling and attenuates α-Synuclein pathology in a Parkinson’s disease mouse model"

**Supplementary Information**

Supplementary file 1. DEG data from RNA-seq, on primary astrocyte of WT and *Acsbg1* KO, treated with PBS, or TNF-α / IL-1α (excel file)

Supplementary file 2. Lipid LC-MS data from conditioned medium, from primary astrocyte of WT and *Acsbg1* KO, treated with PBS, or TNF-α / IL-1α (excel file)

Supplementary table 1. Materials used for this study

| **Reagent or resource** | **Source** | **Identifier** |
|---|---|---|
| **Antibodies** | | |
| Anti-Alpha-synuclein antibody (clone 42) | BD | RRID: AB_398107 |
| p-S129 Synuclein (clone EP1536Y) | Abcam | RRID: AB_869973 |
| p-S129 Synuclein (clone D1R1R) | Cell Signaling Technology | RRID: AB_279886 |
| Anti-Vinculin | Sigma-Aldrich | RRID: AB_477629 |
| Anti-GFAP | Novus Biologicals | RRID: AB_829022 |
| Anti-Iba1 | Wako Chemicals | RRID: AB_839504 |
| Anti-Tyrosine Hydroxylase (TH) | EMD Millipore | RRID: AB_390204 |
| Anti-Acsbg1 | Atlas Antibodies | RRID:AB_2677593 |
| Alexa Fluor® 488 AffiniPure Donkey Anti-Mouse IgG (H+L) | Jackson ImmunoResearch | RRID: AB_2340846 |
| Donkey anti-Rabbit IgG (H+L) Highly Cross-Adsorbed Secondary Antibody, Alexa Fluor™ 555 | Thermo Fisher Scientific | RRID: AB_162543 |
| Donkey anti-Goat IgG (H+L) Cross-Adsorbed Secondary Antibody, Alexa Fluor™ 488 | Thermo Fisher Scientific | RRID: AB_2534102 |
| **Cytokine and lipids** | | |
| TNF-alpha Recombinant Protein | Sigma-Aldrich | Catalog# SRP3177 |
| IL-1alpha Recombinant Protein | Sigma-Aldrich | Catalog# SRP3310 |
| Human CCL5 (RANTES) Recombinant Protein | Thermo Fisher Scientific | Catalog# 300-06 |
| Human CCL20 (MIP-3 alpha) Recombinant Protein | Thermo Fisher Scientific | Catalog# 300-29 |
| Oleic Acid | Sigma-Aldrich | Catalog# O3008 |
| Sphingosine (C16:1, C18:1, C20:1, C22;1) | From the Lab | - |
| **Kit** | | |
| VECTASTAIN(R) ELITE(R) ABC anti-Rabbit IgG HRP Immunodetection Kit | Vector Laboratories | Catalog #PK-6101 |
| Mouse Cytokine Antibody Array (Membrane, 62 Targets) | abcam | Catalog # ab133995 |
| Papain dissociation kit | Worthington Biochemical | Catalog # LK003178 |
| **Experimental models: Organisms/strain** | | |
| Mouse: wild type (WT: C57BL/6J) | Jackson Laboratory | Catalog #000664 |
| Mouse: *Thy1-α-Syn TG Line 61* | Jackson Laboratory | RRID: IMSR_JAX:038796 |
| Mouse: *Acsbg1 KO* | Jackson Laboratory | RRID: IMSR_JAX:024754 |
| **Plasmid DNA** | | |
| pcDNA3.1-Acsbg1-(k)DYK | Genescript | Catalogue: OHu06859 |
| pcDNA3.1-Acsbg1$^{K701A}$-(k)DYK | From the Lab | - |
| **Oligonucleotides** | | |

| | | |
|---|---|---|
| Genotyping primer for WT sequence of *Acsbg1 KO* forward: anneal at 58 degrees<br>5'- TCCACAGACAGGTGACTAA -3' | Sigma-Aldrich | N/A |
| Genotyping primer for WT and *Acsbg1 KO* common reverse: anneal at 58 degrees<br>5'- CACTGAACATGGTTTGAGC -3' | Sigma-Aldrich | N/A |
| Genotyping primer for KO sequence of *Acsbg1 KO* forward: anneals at 58 degrees<br>5'- AGTACTGTGGTTTCCCAAATGTGTCA -3' | Sigma-Aldrich | N/A |
| Genotyping primer for Thy1-α-Syn TG reverse: anneals at 57 degrees<br>5'- GATGATGGCATGCAGCACTGG -3' | Sigma-Aldrich | N/A |
| Genotyping primer for Thy1-α-Syn TG reverse: anneals at 57 degrees<br>5'- GACGGGTGTGACAGCAGTAGC -3' | Sigma-Aldrich | N/A |
| Cloning forward primer for Acsbg1$^{K701A}$ mutant: anneals at 58 degrees<br>5'- tcccacgatggcgctgaaacggc -3' | Sigma-Aldrich | N/A |
| Cloning reverse primer for Acsbg1$^{K701A}$ mutant: anneals at 58 degrees<br>5'- cccaactctccaccc -3' | Sigma-Aldrich | N/A |

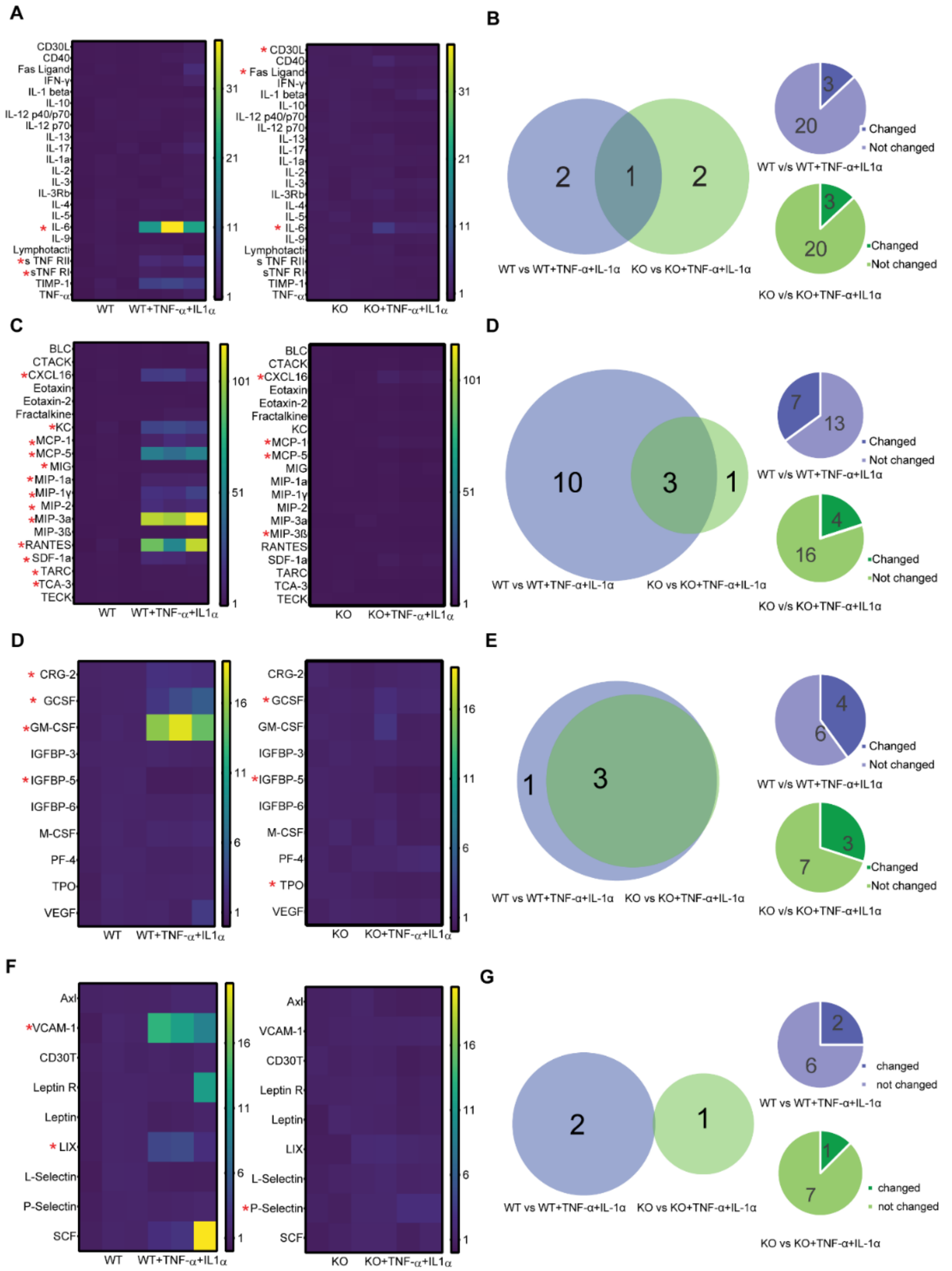


**Supplementary Figure 1. Conditioned media from *Acsbg1* KO show reduced inflammatory ligands secretion from *Acsbg1* KO astrocytes**

(**A, C, D, F**) Heatmaps depicting the levels of cytokines (**A**), chemokines (**C**), growth factors (**D**), and other inflammatory mediators (**F**) measured in conditioned media from wild-type (WT) or Acsbg1 knockout (KO) astrocytes under basal conditions (PBS-treated, naive) or following pro-inflammatory stimulation with TNF-α and IL-1α (pathologically reactive state). Color scale represents relative expression intensity. Analytes marked with a red asterisk (*) indicate statistically significant differences. n=3 per group; *p<0.05.

(**B, D, E, G**) Venn diagrams (**left panels**) illustrating the overlap in significantly changed inflammatory mediators between WT vs. WT+TNF-α+IL-1α (blue) and KO vs. KO+TNF-α+IL-1α (green) comparisons. Pie charts (**right panels**) show the number of cytokines were significantly changed (colored) versus unchanged (light) in WT vs. WT+TNF-α+IL-1α (blue, upper) and KO vs. KO+TNF-α+IL-1α (green, lower).

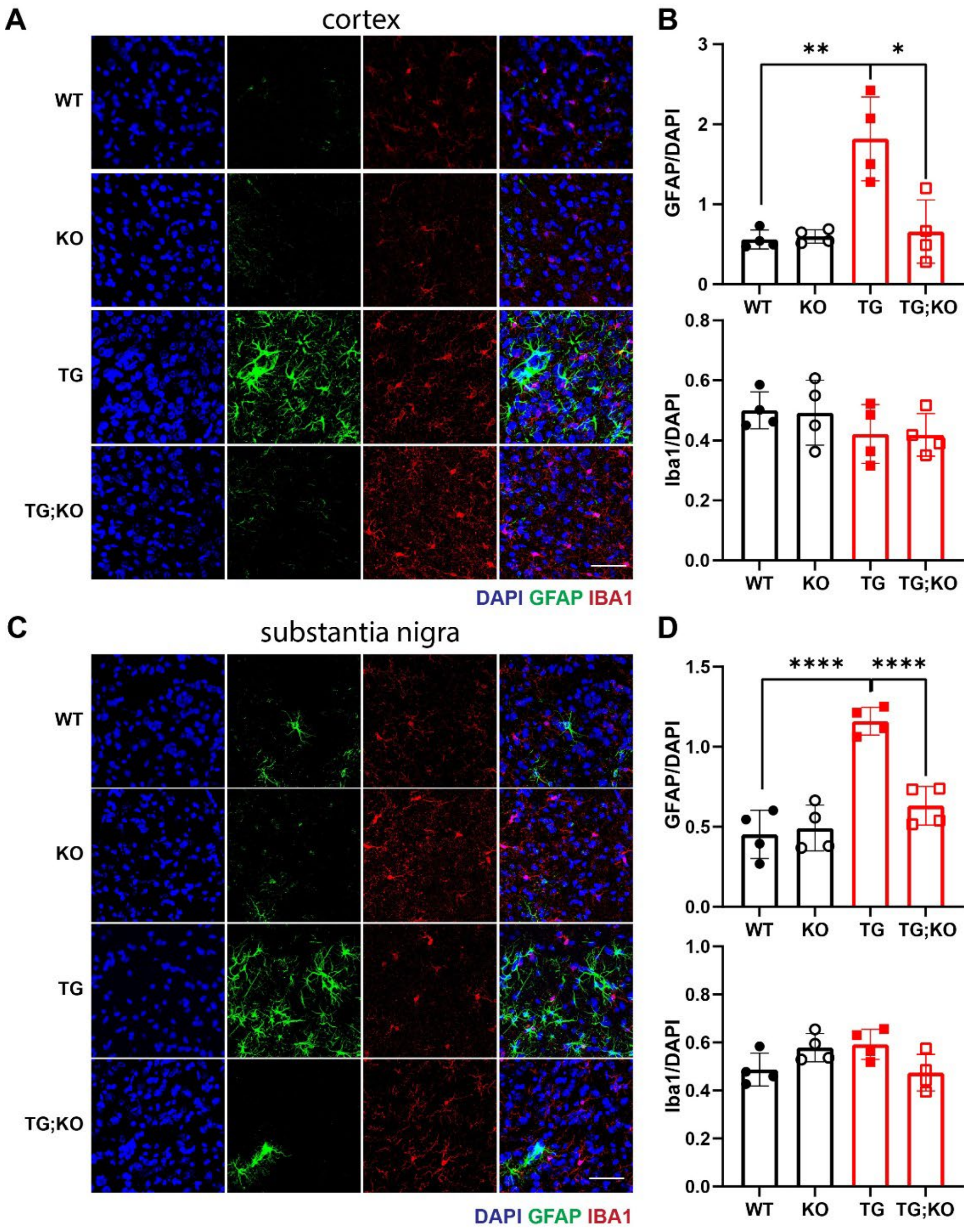


**Supplementary Figure 2.** ***Acsbg1*** **deletion reduces reactive astrogliosis but not microglial activation in Thy1-α-Syn transgenic mice.**

**(A, C)** Representative immunofluorescence images of brain sections stained for DAPI (blue), GFAP (green), and IBA1 (red) from WT, KO, TG, and TG;KO mice at cortex and substantia nigra. Scale bar, 50 µm. **(B)** Quantification of GFAP (top) and IBA1 (bottom) immunoreactivity normalized to DAPI in the regions shown in (A). p* < 0.05, p**<0.01, n = 4 per group. **(D)** Quantification of GFAP (top) and IBA1 (bottom) immunoreactivity normalized to DAPI in the regions shown in (C). **** p < 0.0001, n = 4 per group.

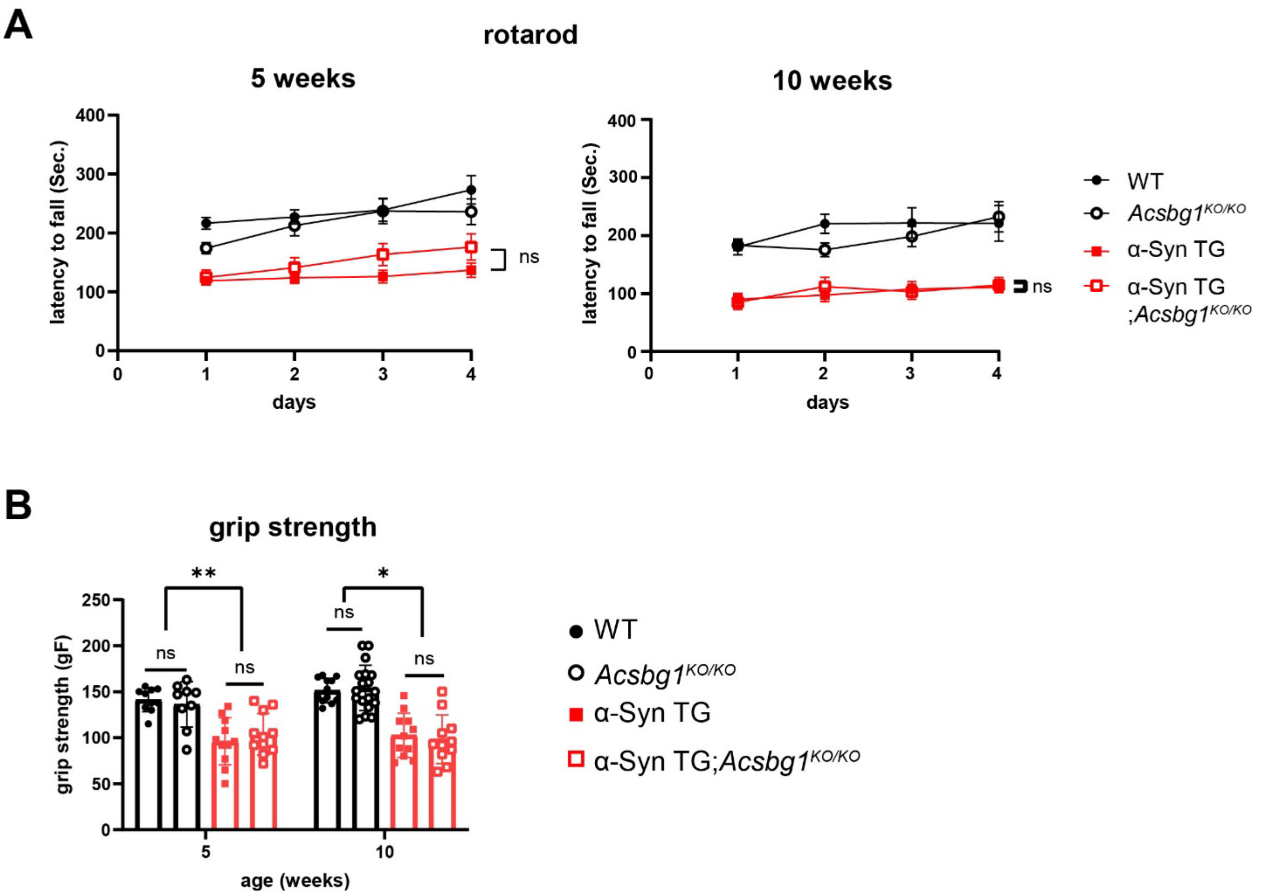


**Supplementary Figure 3. Characterization of the motor function and weight difference of α-syn TG mice with *Acsbg1* KO background.** We compared the motor function of Thy1-α-Syn TG (TG) and its *Acsbg1* KO background (TG;KO) to wild-type and *Acsbg1* KO littermates at 5 and 10 weeks of age. (**A**) motor coordination in the rotarod test, and (**B**) motor strength by grip strength test. * $p<0.05$, ** $p<0.01$, ns=non-significant. WT (n=7 for 5 weeks and 14 for 10 weeks), *Acsbg1*$^{KO/KO}$ (n=15 for 5 weeks, n=16 for 10 weeks), Thy1-α-Syn TG (n=9 for 5 weeks, n=15 for 10 weeks), Thy1-α-Syn TG*; Acsbg1*$^{KO/KO}$ (n=12 for 5 weeks, n=13 for 10 weeks). All mice used for the behavior test are male mice due to Thy1-α-syn TG construct being on the X chromosome.
